# Deep Learning to Predict Future Cognitive Decline: A Multimodal Approach Using Brain MRI and Clinical Data

**DOI:** 10.1101/2025.02.21.639520

**Authors:** Tamoghna Chattopadhyay, Pavithra Senthilkumar, Rahul H. Ankarath, Christopher Patterson, Emma J. Gleave, Sophia I. Thomopoulos, Heng Huang, Li Shen, Lei You, Degui Zhi, Paul M. Thompson

## Abstract

Predicting the trajectory of clinical decline in aging individuals is a pressing challenge, especially for people with mild cognitive impairment, Alzheimer’s disease, Parkinson’s disease, or vascular dementia. Accurate predictions can guide treatment decisions, identify risk factors, and optimize clinical trials. In this study, we compared two deep learning approaches for forecasting changes, over a 2-year interval, in the Clinical Dementia Rating scale ‘sum of boxes’ score (sobCDR). This is a key metric in dementia research, and scores range from 0 (no impairment) to 18 (severe impairment). To predict decline, we trained a hybrid convolutional neural network that integrates 3D T1-weighted brain MRI scans with tabular clinical and demographic features (including age, sex, body mass index (BMI), and baseline sobCDR). We benchmarked its performance against AutoGluon, an automated multimodal machine learning framework that selects an appropriate neural network architecture. Our results demonstrate the importance of combining image and tabular data in predictive modeling for clinical applications. Deep learning algorithms can fuse image-based brain signatures and tabular clinical data, with potential for personalized prognostics in aging and dementia.

## I. Introduction

Predicting future clinical decline in individuals with mild cognitive impairment (MCI) or preclinical stages of other neurodegenerative diseases is critical for early diagnosis, treatment planning, and therapeutic intervention. Neurodegenerative conditions, including Alzheimer’s disease (AD), Parkinson’s disease, and vascular dementia, affect millions worldwide and impose significant personal, societal, and economic costs. Among individuals diagnosed with MCI, approximately 10-15% progress to dementia annually, a transition marked by a subtle yet measurable worsening of clinical symptoms [1]. The Clinical Dementia Rating (CDR) scale [2], particularly the sum of boxes score (sobCDR), is a widely-used measure of the severity of cognitive and functional impairment– rating impairment on a scale of 0 (none) to 18 (severe). Accurate prediction of future sobCDR scores is essential for assessing disease progression, guiding clinical trial design, and tailoring interventions to patient-specific trajectories.

Predicting clinical decline is challenging, as multiple risk factors affect the onset and progression of neurodegenerative diseases. Cognitive and functional impairments in aging individuals are influenced by genetic, environmental, and lifestyle factors. Abnormal brain changes, observable with neuroimaging, may precede clinical symptoms by years or even decades. Predictive models must account for this broad range of causal or influential factors by integrating multimodal data from neuroimaging, demographics, clinical history, and baseline clinical measures. However, the heterogeneity of these data types poses challenges for many traditional machine learning algorithms, which may need to merge information from structured and unstructured inputs. If successful, accurate prognostic models could also be used to gauge treatment effects in clinical trials. As **Figure 1** shows, recent advancements in therapeutic interventions, such as the anti-amyloid drug, lecanemab [3], can significantly reduce the rate of decline as measured by sobCDR; this randomized clinical trial found a 27% slowing after 18 months. Treatment effects would be easier to identify if statistical models could more accurately identify factors that influence disease trajectories based on baseline clinical and neuroimaging data.

**Fig. 1.**
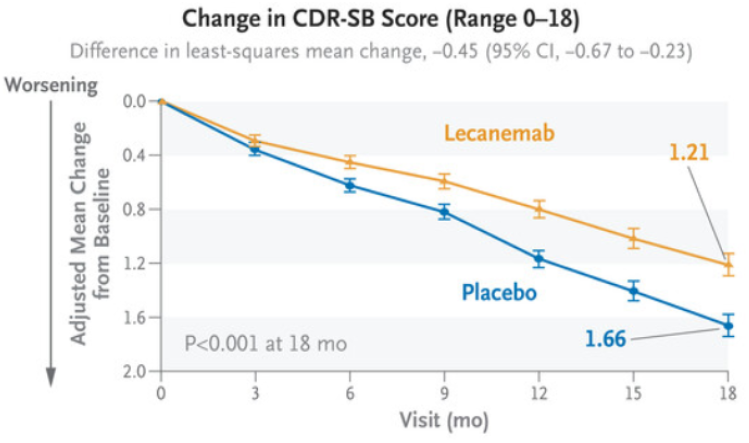
Progressive decline in sobCDR score is a primary outcome measure in many dementia drug trials (here, in a clinical trial evaluating the anti-amyloid drug, lecanemab). Reproduced from van Dyck *et al*. [3]

**Fig. 2.**
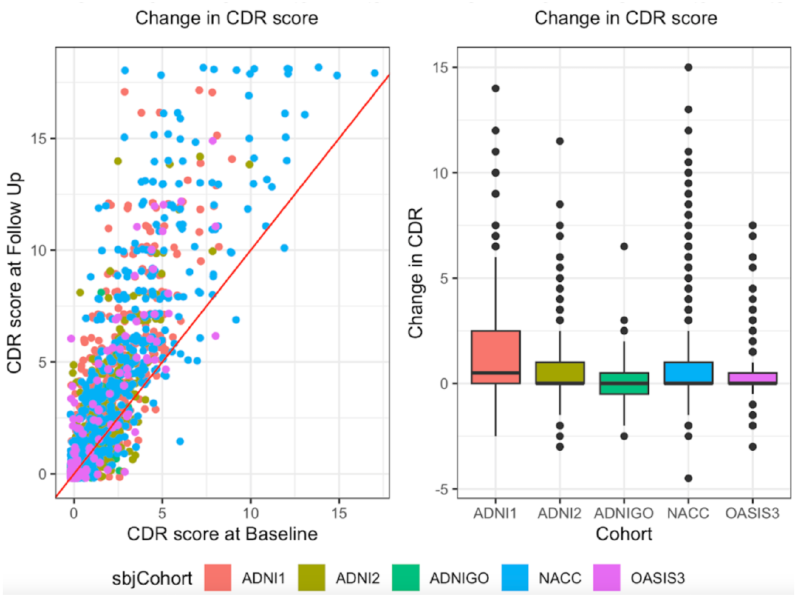
Descriptive statistics summarize the overall trends in the training and test data. Individuals from different cohorts are shown in different colors; the ADNI dataset is divided into 3 components collected during different phases of the study (1, 2, and ‘GO’). Changes in sum-of-boxes score (sobCDR) over 2 years for individual participants in each dataset are shown in the scattergram (*left*) and box-and-whiskers plots (*right*), split by cohort. Higher scores denote greater impairment.

**Fig. 3.**
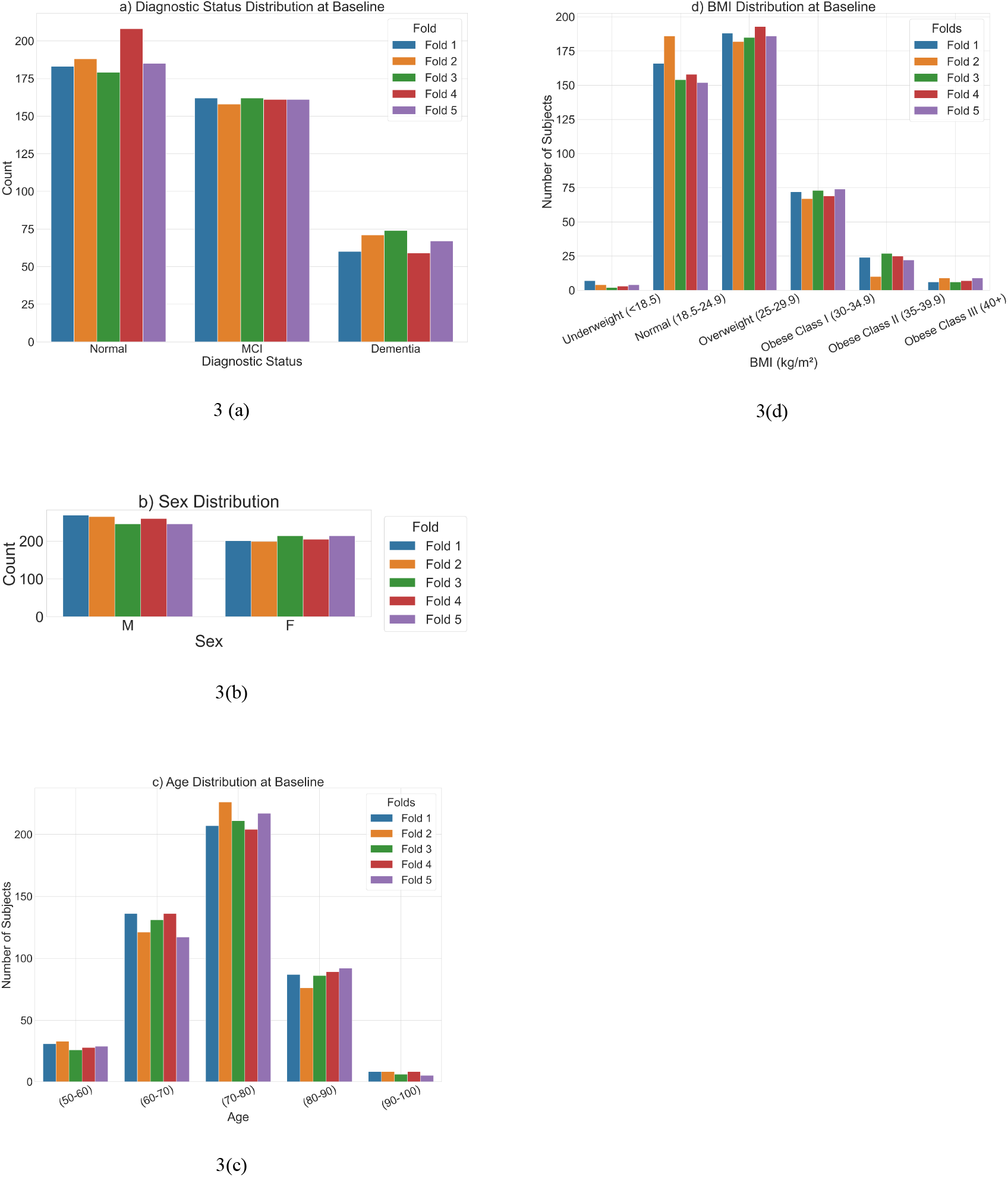
**(a)** Distribution of Diagnostic Status at Baseline, across the five folds used for cross-validation. We also show the distribution of **(b)** Sex, **(c)** age at baseline, and **(d)** BMI at baseline, across the five folds.

Prior work predicting clinical decline from multimodal data includes the TADPOLE challenge [4], in which 33 teams used 92 algorithms to predict AD progression from multimodal biomarkers. Even so, TADPOLE used only one dataset (ADNI), making it unclear how well performance might generalize to other cohorts. Recent work has also leveraged a variety of machine learning and deep learning approaches. Traditional statistical models, such as linear regression [5,6] and Cox proportional hazards models [7,8,9], can be used to predict clinical decline but are not well-suited to including entire images as inputs, or fusing multimodal data streams. More recent innovations include ‘Longitudinal-to-Cross-sectional’ (L2C) transformations to convert variable longitudinal histories into fixed-length feature vectors; other researchers have fitted models to entire longitudinal histories, as in the ‘AD Course Map’ (AD-Map) approaches and methods using minimal recurrent neural networks (MinimalRNN) [10]. More recently, computer vision-based deep learning architectures have been adapted for biomedical image analysis to extract diagnostic or prognostic features, and identify biomarkers of neurodegenerative diseases. For example, convolutional neural networks (CNNs) and vision transformers have been applied to analyze 3D T1-weighted MRI scans, yielding insights into structural brain changes associated with AD [11,12]. AutoML platforms, such as AutoGluon [13], have also gained traction as they can optimize and ensemble models across diverse datasets, or select neural network architectures that perform best for a task–achieving competitive performance with minimal manual intervention.

In this paper, we compare our custom hybrid deep learning model – that integrates tabular and image data – with the AutoGluon framework, on the task of predicting two-year changes in sobCDR scores. Our hybrid model uses a CNN to extract meaningful features from 3D T1-weighted brain MRI scans and combines these with fully connected layers that process tabular inputs, including demographic and clinical data. AutoGluon [13]–an automated machine learning framework–serves as a robust baseline for comparison. The contributions of this work are two-fold: (1) we show the feasibility and advantages of a hybrid deep learning approach to integrate multimodal data, and (2) we benchmark the performance of an AutoML platform against our custom-designed architecture. By addressing these aspects, our work lays the groundwork for improving prognostic modeling in neurodegenerative diseases and fostering the development of personalized predictive models.

## II. Data and Preprocessing

To evaluate the proposed predictive models, we used data from three independent, publicly available cohorts: (1) the Alzheimer’s Disease Neuroimaging Initiative [14] (ADNI; 1,136 participants: 73.72±7.11 years, 507 F/629 M; 397 cognitively normal (CN), 593 MCI, 146 AD), (2) the Open Access Series of Imaging Studies [15] (OASIS-3; 241 participants: 72.88±6.44 years, 113 F/128 M; CN/MCI/AD), and (3) the National Alzheimer’s Coordinating Center [16] (NACC; 942 participants: 72.23 ± 9.38 years, 413 F/529 M; 546 CN, 211 MCI, 185 AD). We included participants who met the following inclusion criteria: availability of a 3D T1-weighted brain MRI at baseline, baseline measurements of age, sex, and body mass index (BMI), a baseline sum of boxes Clinical Dementia Rating (sobCDR) score recorded within 90 days of the imaging session, and a follow-up sobCDR score measured between 1.75 and 2.25 years after the baseline visit. These criteria ensured that all participants had comparable data for longitudinal analysis of clinical decline.

For evaluation, datasets were partitioned into five independent cross-validation folds [17], stratified to ensure balanced distributions of sex, baseline age at imaging, and changes in sobCDR scores. This stratification was crucial to mitigate potential biases and to ensure that each fold represented a representative subset of the data, with changes in sobCDR ranging from 0 (no impairment) to 18 (severe impairment).

A preprocessing pipeline was applied to all imaging data to standardize and enhance the quality of the input features. T1-weighted 3D brain MRI volumes were pre-processed using the following steps: nonparametric intensity normalization (N4 bias field correction), ‘skull stripping’ for brain extraction, nonlinear registration to a template with 6 degrees of freedom and isometric voxel resampling to 2 mm [18,19]. Pre-processed images were of size 91×109×91. The T1w images were scaled, via min-max scaling, to the range 0 and 1. All T1w images were aligned to a common template provided by the ENIGMA consortium [20].

## III. Deep Learning Architectures

Our proposed hybrid model [21] fuses a 3D convolutional neural network (CNN) with an artificial neural network (ANN), in a Y-shaped architecture (**Fig. 4**) to combine imaging and tabular data for predicting clinical decline. The 3D CNN processes 3D volumetric T1-weighted MRI scans, while the ANN ingests discrete tabular features, including age, sex, BMI, and baseline sobCDR. These two data modalities are initially processed independently, before being fused for final prediction. The network is trained end to end.

**Fig. 4.**
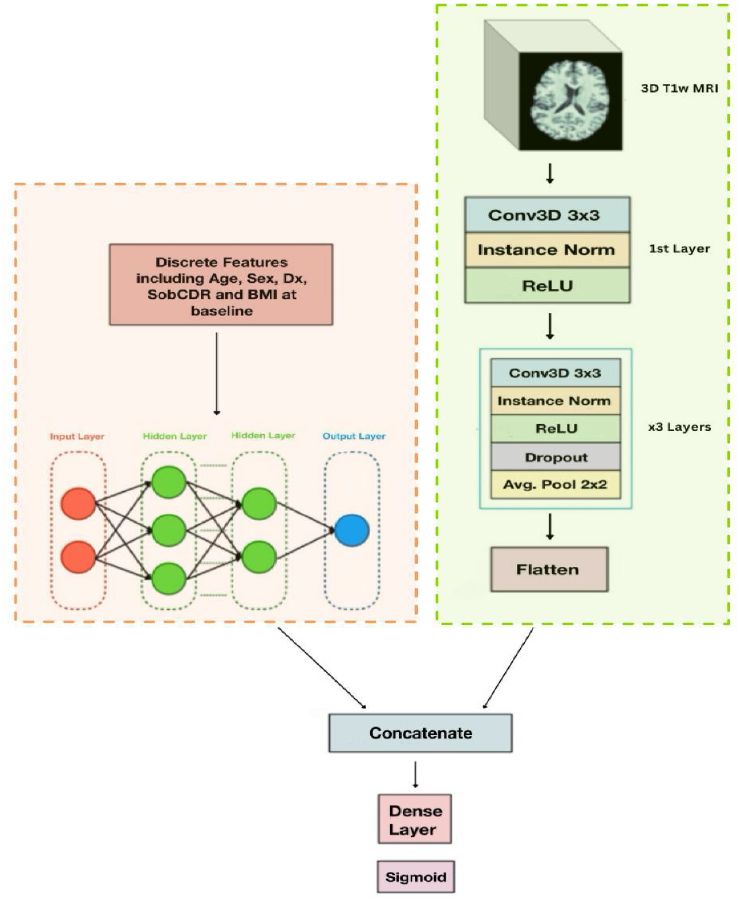
Hybrid 3D CNN Model Architecture. The Y-shaped design merges discrete features in tabular format (left) with entire 3D images (right), which are handled with a 3D CNN. The entire model is trained end-to-end.

**Fig. 5.**
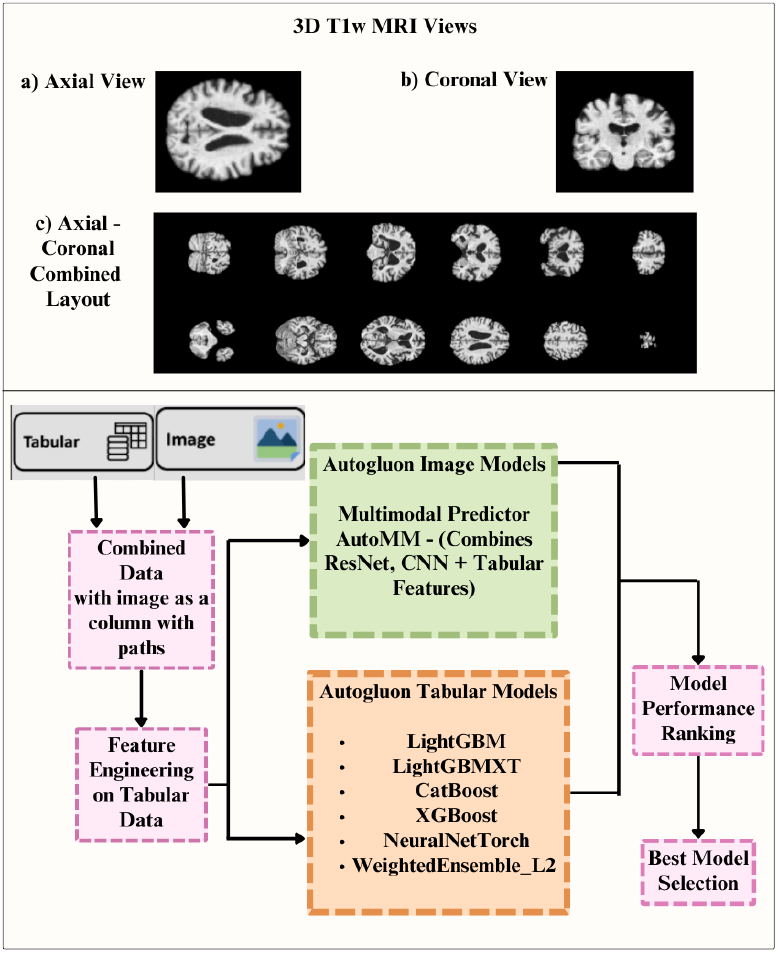
**Top:** Image inputs for the AutoGluon models. **Bottom:** AutoGluon architecture, which automatically selects a neural network architecture from a range of traditional models.

**Fig. 6.**
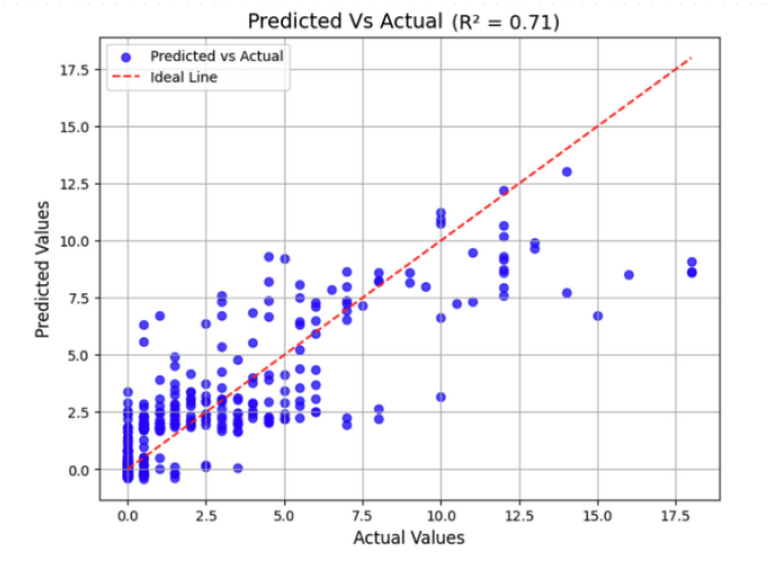
Actual vs predicted Scores (R^2^ score) when tested on fold 3 using the hybrid CNN architecture.

**Fig. 7.**
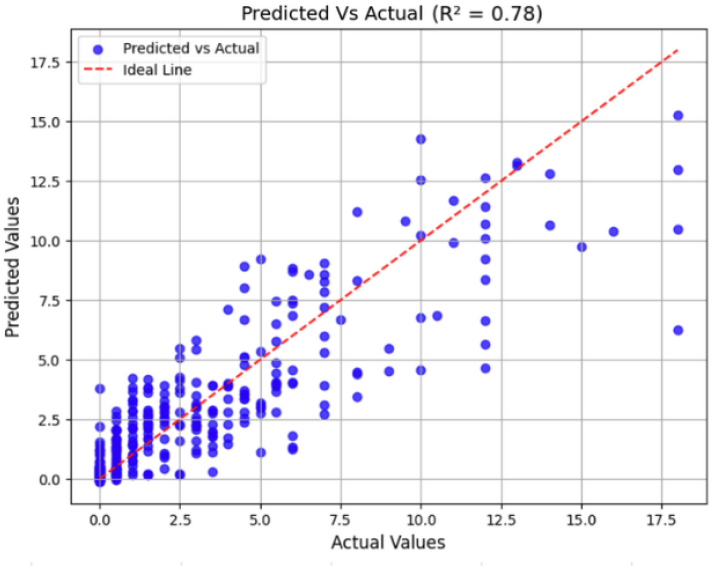
Actual vs predicted Scores (R^2^ score) when tested on fold 3 using AutoGluon for axial middle slice with Standard Scaled Features and R^2^ validation.

The 3D CNN component has three convolutional blocks, each incorporating filters of increasing size (32, 64, 128, and 256) to progressively extract hierarchical features from the MRI scans. Each convolutional block is followed by Batch Normalization and Max Pooling layers to enhance training stability and spatial feature reduction. The final convolutional layer, before concatenation, uses a filter size of 256 and employs average pooling to retain salient global spatial features while minimizing dimensionality. In parallel, the ANN component processes the tabular data using three fully connected layers with hidden sizes of 1024, 512, and 64, respectively. Each layer is accompanied by instance normalization and the ReLU activation function to ensure stable and efficient learning. The structured numerical and categorical variables are first normalized and embedded appropriately, before being passed through the ANN.

After processing through their respective architectures, the feature representations from the CNN and ANN components are concatenated via tensor stacking. The combined feature map is then passed through additional fully connected Dense layers to produce the final prediction of the sobCDR score at two years. The model was trained with a learning rate of 0.001 using the Adam optimizer, with a batch size of 2 to handle the computational demands of 3D imaging data. Training was conducted over 200 epochs, ensuring sufficient convergence, by stopping training when the validation loss did not decrease further over 10 epochs, while preventing overfitting. The model’s generalizability was validated using five-fold cross-validation, where the hybrid model was independently trained on four folds, and tested on the held out fifth sector of the data.

Performance was assessed using mean absolute error (MAE) and the coefficient of determination (R^2^ score). The MAE and R^2^ values were recorded for each cross-validation fold, with the final performance determined as the average of these metrics across all five folds. This ensured robust assessment of the model’s ability to predict longitudinal cognitive decline.

To provide a comparative benchmark, we used AutoGluon [13], an automated machine learning (AutoML) framework, which automatically selects a neural network architecture from a diverse set of traditional models, and trains it to predict future sobCDR scores. AutoGluon is well-suited for multimodal learning tasks, as it supports tabular, text, and image-based inputs with minimal user intervention. For the image input, rather than using full 3D brain MRIs, we extracted the 2D axial and coronal middle slices from the T1-weighted brain images. This approach reduces computational complexity while preserving structural information from relevant regions of the brain. The same discrete variables used for our hybrid deep learning model—age, sex, BMI, and baseline sobCDR—were included in the tabular input. AutoGluon automatically pre-processes the data, selects an appropriate neural network architecture, and tunes hyperparameters to optimize predictive performance. The training process involved automated selection of model ensembling strategies, ensuring robust predictions while reducing the risk of overfitting. We evaluated AutoGluon’s performance using the same five-fold cross-validation approach as our hybrid model, calculating average MAE and R^2^ scores across the folds.

## IV. Experimental Results

Our hybrid deep learning model was evaluated using five-fold cross-validation, with performance metrics recorded for each fold. The mean absolute error (MAE)–which measures the average deviation of predicted sobCDR scores from actual values–yielded an average of 1.303 across all folds, indicating strong predictive accuracy. In other words, the follow-up score is predicted with a mean error of just over 1 point, on an 18 point scale. The model’s coefficient of determination (R^2^), which quantifies the proportion of variance explained by the model, achieved an average of 0.704, demonstrating a strong correlation between predicted and actual clinical decline. Performance was consistent across folds: MAE values ranged from 1.231 to 1.414, and R^2^ values ranged from 0.659 to 0.763, highlighting the model’s robustness and reliability in forecasting cognitive decline over a two-year period.

We evaluated the performance of AutoGluon’s *MultimodalFusionMLP* model in predicting clinical decline using a combination of T1-weighted MRI middle axial slices and demographic features. The model was trained with approximately 97.4M parameters using two different validation approaches and two scaling methods, for the tabular features, and assessed through 5-fold cross-validation. Using standard scaling, the model achieved an average R^2^ score of 0.763 and MAE of 1.002 when optimized for MSE validation. When optimized for R^2^ validation, the model showed similar performance with an average R^2^ of 0.771 and MAE of 0.968. With MinMax scaling, the model demonstrated comparable results. Under MSE validation, it achieved an average R^2^ score of 0.761 and MAE of 1.022. The R^2^ validation configuration yielded an average R^2^ of 0.757 and MAE of 1.011. Across all configurations, the model maintained consistent performance, with R^2^ scores ranging from 0.735 to 0.808, indicating robust predictive capability. The MAE values were stable across folds, typically between 0.94 and 1.08, suggesting reliable prediction accuracy regardless of the validation metric or scaling method chosen.

The performance of AutoGluon’s *MultimodalFusionMLP* model when using a combination of T1-weighted MRI middle coronal slices and demographic features was slightly better. The same training configurations were used as the previous case - two different validation approaches and two different scaling methods, assessed through 5-fold cross validation. Using standard scaling, the model achieved an average R^2^ score of 0.759 and MAE of 0.983 across both validation models. Using min-max scaling, the model achieved an average R^2^ score of 0.749 and MAE of 0.988 across both validation models.

As an ablation study, a simple general linear model was constructed to estimate the predictive capacity when using only the tabular covariates to estimate the outcome across the five folds. This model does not take into account any imaging information at all. Comparing the results to AutoGluon, it shows that adding imaging features helps to improve the accuracy of the prediction.

**TABLE 1.** Performance of deep learning architectures for predicting future clinical decline. For the AutoGluon models, *axial* means that the model used the middle axial slice as the image input, *coronal* means it used the middle coronal slice, and AC means it used multiple axial and coronal slices as the image input. *s* denotes standard scaling, *m* indicates MinMax scaling, *M* indicates MSE as the validation metric and *R* (used here as a suffix) indicates R^2^ as the validation metric. Best results are in **bold**.

| Model | Metric | Fold 1 | Fold 2 | Fold 3 | Fold 4 | Fold 5 | Avg |
| --- | --- | --- | --- | --- | --- | --- | --- |
| *Linear Null Model* | MAE | 0.980 | 1.027 | 1.13 | 0.998 | 1.012 | 1.096 |
| | $R^2$ | 0.786 | 0.804 | 0.757 | 0.740 | 0.744 | 0.767 |
| *Hybrid CNN* | MAE | 1.266 | 1.263 | 1.342 | 1.231 | 1.413 | 1.303 |
| | $R^2$ | 0.763 | 0.658 | 0.694 | 0.716 | 0.687 | 0.704 |
| *AutoGluon axial sM* | MAE | 0.946 | 0.964 | 1.083 | 1.057 | 0.958 | 1.002 |
| | $R^2$ | 0.808 | 0.761 | 0.735 | 0.753 | 0.760 | 0.763 |
| *AutoGluon axial sR* | MAE | 0.957 | 0.945 | 0.991 | 0.967 | 0.977 | **0.968** |
| | $R^2$ | 0.794 | 0.767 | 0.745 | 0.777 | 0.772 | **0.771** |
| *AutoGluon axial mM* | MAE | 1.048 | 1.072 | 1.051 | 0.964 | 0.976 | 1.022 |
| | $R^2$ | 0.788 | 0.740 | 0.744 | 0.774 | 0.759 | 0.761 |
| *AutoGluon axial mR* | MAE | 1.032 | 1.048 | 1.032 | 0.968 | 0.975 | 1.011 |
| | $R^2$ | 0.754 | 0.753 | 0.750 | 0.765 | 0.761 | 0.757 |
| *AutoGluon coronal sM* | MAE | 0.975 | 0.919 | 1.002 | 1.002 | 1.018 | 0.983 |
| | $R^2$ | 0.787 | 0.768 | 0.726 | 0.752 | 0.760 | 0.759 |
| *AutoGluon coronal sR* | MAE | 0.975 | 0.919 | 1.002 | 1.002 | 1.018 | 0.983 |
| | $R^2$ | 0.787 | 0.768 | 0.726 | 0.752 | 0.760 | 0.759 |
| *AutoGluon coronal mM* | MAE | 0.887 | 1.052 | 1.059 | 0.949 | 0.995 | 0.988 |
| | $R^2$ | 0.793 | 0.728 | 0.722 | 0.757 | 0.744 | 0.748 |
| *AutoGluon coronal mR* | MAE | 0.887 | 1.052 | 1.059 | 0.949 | 0.995 | 0.988 |
| | $R^2$ | 0.793 | 0.728 | 0.722 | 0.757 | 0.744 | 0.748 |
| *AutoGluon AC sM* | MAE | 0.976 | 0.902 | 1.033 | 1.021 | 1.028 | 0.992 |
| | $R^2$ | 0.789 | 0.778 | 0.749 | 0.763 | 0.772 | 0.771 |
| *AutoGluon AC sR* | MAE | 0.941 | 0.968 | 0.992 | 1.017 | 0.986 | 0.981 |
| | $R^2$ | 0.804 | 0.778 | 0.739 | 0.756 | 0.784 | 0.773 |
| *AutoGluon AC mM* | MAE | 1.034 | 0.897 | 1.026 | 1.015 | 1.022 | 0.999 |
| | $R^2$ | 0.728 | 0.779 | 0.750 | 0.764 | 0.774 | 0.759 |
| *AutoGluon AC mR* | MAE | 0.996 | 0.961 | 0.987 | 1.012 | 0.981 | 0.987 |
| | $R^2$ | 0.750 | 0.779 | 0.741 | 0.758 | 0.785 | 0.763 |

The performance of AutoGluon’s *MultimodalFusionMLP* model did not greatly improve when using a combination of multiple T1-weighted MRI middle coronal and axial slices and demographic features. The same training configurations were used as the previous case - two different validation approaches and two different scaling methods, assessed through 5-fold cross validation. Using standard scaling, the model achieved an average R^2^ score of 0.773 and MAE of 0.981. Using min-max scaling, the model achieved an average R^2^ score of 0.759 and MAE of 0.999. These results are consistent with those when only a single slice was used as the image input.

## V. Discussion

Our hybrid deep learning model demonstrated strong predictive performance in forecasting sobCDR progression, as evidenced in the 5-fold cross-validation. The model achieved an average MAE of 1.303, indicating high accuracy in estimating cognitive decline, with consistent performance across folds (MAE range: 1.231–1.414). Additionally, the R^2^ score averaged 0.704, reflecting a strong correlation between predicted and actual clinical scores. These results highlight the model’s robustness and reliability across multiple cohorts and datasets, suggesting its promise for real-world clinical applications. Minor variations were observed across folds, but the model consistently captured predictors needed to forecast disease progression.

AutoGluon’s *MultimodalFusionMLP* model performed well for predicting clinical decline using neuroimaging and demographic data. The consistent R^2^ values around 0.75-0.77 across different validation approaches and scaling methods suggest that the model captures around 76% of the variance in clinical outcomes, which is promising for this complex medical outcome. Performance did not further improve when using multiple slices versus using a single brain MRI slice as the image input. Several key observations emerge from our analysis:

- **Scaling Method:** Standard scaling and MinMax scaling produced comparable results, with standard scaling yielding marginally better performance (R^2^ = 0.771 vs 0.757 under R^2^ validation). This suggests that the model is robust to the choice of feature scaling method, though standard scaling might offer a slight advantage in capturing the underlying relationships in the data.
- **Validation Metric:** The similar performance on MSE and R^2^ validation metrics (MAE ≈ 1.0 across configurations) indicates that the models’ predictive capability is stable regardless of the optimization criterion. This robustness is valuable in clinical applications where consistency across different evaluation methods is crucial.
- **Cross-validation Stability:** The relatively small variations in performance across folds (R^2^ standard deviation ≈ 0.013-0.027) suggest that the model generalizes well across different subsets of the data. This stability is needed for clinical applications, where reliable performance across diverse patient populations is essential.
- **Clinical Implications:** The achieved MAE of around 1 point in predicting future decline represents a clinically meaningful level of accuracy, considering the scale and complexity of the factors that affect future clinical decline. This level of accuracy may potentially support clinical decision making. Realistically, the lower bound on prediction error depends on the reliability of the clinical test itself, as the test does not have perfect test-retest reliability when administered by different raters, or on different days. Prior work on the sobCDR score suggests that test-retest reliability is good (intra-class correlation coefficient [ICC]=0.83) [22]; in one study, 24 human raters independently scored the CDR using four videotaped interviews [23], and there was moderate to high overall interrater reliability (*kappa* statistic: 0.62), despite some difficulties in reliably assessing early dementia. Even so, a more recent study [24] of 139 patients (age: 80.1 ± 6SD, 72% with dementia) reported a much higher ICC of 0.95 (95% CI: 0.93–0.97) for the sobCDR score assessed face-to-face and with all the information available in the patient’s medical record. The mean difference between the sobCDR score assessed face-to-face and with the medical record was 0.098 ± 1.036. In any case, ICC statistics would be valuable to ascertain for the raters in future benchmarking studies, as the clinical test inter-rater and test-retest reliability essentially define the achievable limits of machine learning model performance.

## VI. Conclusion and Future Work

Our results show the potential of multimodal deep learning for accurately forecasting cognitive decline–a line of work that could aid in early diagnosis and personalized treatment planning for neurodegenerative diseases. While the models showed robustness across different validation folds, future work will attempt to further improve accuracy by incorporating additional data types, potentially including diffusion MRI or PET data, genome-wide genetics, and blood-derived proteomic markers, as well as longitudinal clinical assessments. Additionally, advanced techniques such as self-supervised learning and transformer-based architectures that can handle missing data [25] may further enhance performance, and may also help to predict decline in specific subdomains of cognitive performance. Finally, validating the model on new and more diverse clinical datasets will be crucial for assessing real-world applicability and potential integration into clinical decision-making frameworks.

## Acknowledgments

This work was supported by NIH NIA grants U01AG068057 (‘AI4AD’), U01 AG066833, and U01AG070112.

## References

[1] World Health Organization, “Dementia,” 2022. https://www.who.int/news-room/fact-sheets/detail/dementia.

[2] Cedarbaum, J. M., et al. (2013). ‘Rationale for use of the Clinical Dementia Rating Sum of Boxes as a primary outcome measure for Alzheimer’s disease clinical trials.’ Alzheimer’s & Dementia, 9(1), S45–S55.

[3] Van Dyck, C. H., et al. (2023). ‘Lecanemab in early Alzheimer’s disease.’ New England Journal of Medicine, 388(1), 9–21.

[4] Marinescu, R. V., et al. (2019). TADPOLE Challenge: Accurate Alzheimer’s disease prediction through crowdsourced forecasting of future data. In Predictive Intelligence in Medicine: Second International Workshop, PRIME 2019, Held in Conjunction with MICCAI 2019, Shenzhen, China, October 13, 2019, Proceedings 2 (pp. 1–10). Springer International Publishing.

[5] Dickerson, B. C., et al. (2007). Clinical prediction of Alzheimer disease dementia across the spectrum of mild cognitive impairment. Archives of General Psychiatry, 64(12), 1443–1450.

[6] Mattsson-Carlgren, N., et al. (2023). Prediction of longitudinal cognitive decline in preclinical Alzheimer disease using plasma biomarkers. JAMA Neurology, 80(4), 360–369.

[7] Wang, M., et al. (2022). Dementia risk prediction in individuals with mild cognitive impairment: a comparison of Cox regression and machine learning models. BMC Medical Research Methodology, 22(1), 284.

[8] Rongve, A., et al. (2016). Cognitive decline in dementia with Lewy bodies: a 5-year prospective cohort study. BMJ Open, 6(2), e010357.

[9] Kim, W. J., et al. (2019). Cox proportional hazard regression versus a deep learning algorithm in the prediction of dementia: an analysis based on periodic health examination. JMIR Medical Informatics, 7(3), e13139.

[10] Zhang, C., et al. (2024). Cross-dataset Evaluation of Dementia Longitudinal Progression Prediction Models. medRxiv, 2024–11.

[11] Chattopadhyay, T., et al. (2024, November). Comparison of Explainable AI Models for MRI-Based Alzheimer’s Disease Classification. In 2024 20th International Symposium on Medical Information Processing and Analysis (SIPAIM) (pp. 1–4). IEEE.

[12] Chattopadhyay, T., et al. “Brain Age Analysis and Dementia Classification using Convolutional Neural Networks trained on Diffusion MRI: Tests in Indian and North American Cohorts,” 2024 46th Annual International Conference of the IEEE Engineering in Medicine and Biology Society (EMBC), Orlando, FL, USA, 2024, pp. 1–7.

[13] Erickson, N., et al. (2020). Autogluon-tabular: Robust and accurate autoML for structured data. arXiv preprint 2003.06505.

[14] Jack Jr, C. R., et al. (2008). ‘The Alzheimer’s Disease Neuroimaging Initiative (ADNI): MRI methods.’ Journal of Magnetic Resonance Imaging: An Official Journal of the International Society for Magnetic Resonance in Medicine, 27(4), 685–691.

[15] LaMontagne, P. J., et al. (2019). ‘OASIS-3: longitudinal neuroimaging, clinical, and cognitive dataset for normal aging and Alzheimer disease.’ medrxiv, 2019–12.

[16] Beekly, D. L., et al. (2007). ‘The National Alzheimer’s Coordinating Center (NACC) database: the uniform data set.’ Alzheimer Disease & Associated Disorders, 21(3), 249–258.

[17] Bates, S., Hastie, T., & Tibshirani, R. (2024). Cross-validation: what does it estimate and how well does it do it? J. American Statistical Association, 119(546), 1434–1445.

[18] Chattopadhyay, T., et al. (2023). Predicting dementia severity by merging anatomical and diffusion MRI with deep 3D convolutional neural networks. In the 18th International Symposium on Medical Information Processing and Analysis (SIPAIM; Vol. 12567, pp. 90–99). SPIE.

[19] Lam, P., et al., “3-D Grid-Attention Networks for Interpretable Age and Alzheimer’s Disease Prediction from Structural MRI,” (2020).

[20] Jahanshad, N., et al. (2013). Multi-site genetic analysis of diffusion images and voxelwise heritability analysis: a pilot project of the ENIGMA-DTI working group. NeuroImage, 81, 455–469.

[21] Chattopadhyay, T., et al. (2024). ‘Comparison of deep learning architectures for predicting amyloid positivity in Alzheimer’s disease, mild cognitive impairment, and healthy aging, from T1-weighted brain structural MRI.’ Frontiers in Neuroscience, 18, 1387196.

[22] McDougall, F., et al. (2021). Psychometric properties of the Clinical Dementia Rating—Sum of Boxes and other cognitive and functional outcomes in a prodromal Alzheimer’s disease population. The Journal of Prevention of Alzheimer’s Disease, 8, 151–160.

[23] Rockwood K, et al. Interrater reliability of the Clinical Dementia Rating in a multicenter trial. J Am Geriatr Soc. 2000 May;48(5):558–9.

[24] Dauphinot, V., et al. (2024). Reliability of the assessment of the clinical dementia rating scale from the analysis of medical records in comparison with the reference method. Alzheimer’s research & therapy, 16(1), 198.

[25] Shirkavand, R., et al. (2023). Incomplete Multimodal Learning for Complex Brain Disorders Prediction. arXiv preprint 2305.16222.

[26] Delor, I., et al. (2013). ‘Modeling Alzheimer’s disease progression using disease onset time and disease trajectory concepts applied to CDR-SOB scores from ADNI.’ CPT: Pharmacometrics & Systems Pharmacology, 2(10), 1–10.

[27] O’Bryant, S. E., et al. (2010). ‘Validation of the new interpretive guidelines for the clinical dementia rating scale sum of boxes score in the National Alzheimer’s Coordinating Center database.’ Archives of Neurology, 67(6), 746–749.

[28] Williams, M. M., et al. (2013). ‘Progression of Alzheimer’s disease as measured by Clinical Dementia Rating Sum of Boxes scores.’ Alzheimer’s & Dementia, 9(1), S39–S44.

